# A spatial map of antennal-expressed olfactory ionotropic receptors in the malaria mosquito

**DOI:** 10.1101/2022.05.11.491386

**Authors:** Joshua I. Raji, Christopher J. Potter

## Abstract

The malaria mosquito *Anopheles coluzzii* uses odors to guide various behaviors such as host-seeking. The detection of behaviorally relevant odors is mediated by a diverse family of receptors including the olfactory Ionotropic Receptors (IRs). Olfactory receptors are expressed on olfactory neurons, with the mosquito antennae representing the main olfactory appendage for detecting volatile chemical cues from the environment. It is currently unknown how many neurons across the antenna express a certain IR, or how these IR-expressing neurons are spatially arranged. Here, we performed whole mount fluorescent in situ hybridization of all IRs expressed in the antennae. The organization of IR-positive cells within an antennal segment (flagellomere) appeared stereotyped across multiple antennae. The spatial map of IR-expressing neurons revealed that the antenna might be organized into proximal and distal functional domains. Highly expressed tuning (odor-binding) IRs exhibit distinct co-localization patterns with cognate IR co-receptor(s) in a combinatorial fashion that might predict their functional properties. These findings reveal organizing principles of *Anopheles* IR-expressing neurons in the mosquito which might underlie their functional contribution to the detection of behaviorally relevant odors.

## Introduction

There are ∼3500 species of mosquito on earth, yet the most dangerous are the ∼100 species that serve as vectors of human diseases (Harbach, 2007). Among these species, *Anopheles* and *Aedes* mosquitoes represent the two most important genera of medical health significance to humans. The process of seeking human hosts and blood feeding is a critical conduit for the transmission of pathogens by disease-carrying mosquitoes. Some members of the *Anopheles gambiae* complex, including *An. gambiae* s.s. (S form) and *An. coluzzii* (M form) (Coetzee et al., 2000), are major vectors of malaria, largely because they strongly prefer to host-seek humans (Coutinho-Abreu et al., 2022; Konopka et al., 2021; Raji and DeGennaro, 2017). *Anopheles* rely primarily on their olfactory system to find mates, avoid predators, search for suitable oviposition sites, and locate blood hosts (Konopka et al., 2021). The ability to detect sensory cues is enabled by the possession of sensory receptors expressed by neurons housed in specialized sensillar hairs in the olfactory appendages - the antennae, maxillary palps, and labella.

The mosquito antenna is the principal olfactory organ for detecting odors. The female *Anopheles* antenna contains 3 different types of sensilla categorized based on size and shape identified across 12 antennal segments (flagellomeres) (McIver, 1982). A total of 33 coeloconic sensilla, 84 grooved peg sensilla, and ∼630 trichoid sensilla were identified in the antennae of female *An. gambiae* mosquitoes (McIver, 1982). Additional sensory sensilla have been identified on the maxillary palps and the labella (McIver, 1982; Pitts and Zwiebel, 2006). Each sensillum contains on average between 2-4 neurons that are activated by volatile odorants (Kwon et al., 2006; McIver, 1982). The female *An. gambiae* antenna is densely populated with ∼1500 olfactory receptor neurons, mostly housed in the trichoid sensilla (Pitts and Zwiebel, 2006; Qiu et al., 2006). These neurons express odor-gated sensory receptors that bind odor ligands, ultimately leading to neuronal activation.

Three main classes of odor-gated receptors have been identified in mosquitoes. The Odorant Receptors (ORs), which function with an obligate Odorant Receptor Co-receptor (Orco), are found in antennal trichoid, maxillary palp capitate peg, and labella T2 sensilla. The olfactory Gustatory Receptors (GRs), which function as a complex to detect carbon dioxide, are found on a single neuron in maxillary palp capitate peg sensilla. The olfactory Ionotropic Receptors (IRs), which function with 1 or more IR co-receptors (IR8a, IR25a, IR76b), reside in the antennal grooved peg sensilla, coeloconic sensilla, and possibly trichoid sensilla (Mika and Benton, 2021; Wicher and Miazzi, 2021). Among these three receptor classes, the IRs are the most ancestral with evolutionary history that differs from the ORs and GRs (Benton et al., 2009; Croset et al., 2010).

Functionally, mosquito ORs respond to a variety of plant-derived volatiles and human skin emanations (Carey et al., 2010; DeGennaro et al., 2013; Sun et al., 2020) while olfactory GRs respond primarily to carbon dioxide (McMeniman et al., 2014; Sorrells et al., 2021; Xu et al., 2020). Mosquito IRs are responsive to acids and amines (Obaldia et al., 2022; Raji et al., 2019; Ye et al., 2021), and might be involved in human host preferences (Obaldia et al., 2022). However, the functional roles of mosquito IRs extend beyond sensing behaviorally relevant odorants. Several lines of evidence suggest multiple roles for insect IRs including oviposition cue detection (Chen and Amrein, 2017), courtship behavior (Grosjean et al., 2011), thermosensation (Greppi et al., 2020; Laursen et al., 2022), humidity detection (Knecht et al., 2016; Laursen et al., 2022), and taste sensation (Hussain et al., 2016; Jové et al., 2020). The diverse roles of IRs remain consistent with the widespread expression pattern across insect body parts. In *Drosophila*, IRs are broadly expressed and can be detected in the legs, pharynx, labellum, and wings (Koh et al., 2014). Transcriptomic profiling of *Aedes aegypti* mosquitoes also revealed broad expression of IRs in the olfactory appendages as well as in the legs, rostrum, and abdominal tip (Matthews et al., 2016). Similarly, elevated IR transcripts levels were detected in the body of *Anopheles gambiae* mosquitoes (Pitts et al., 2017).

While the antennal topography of a limited number of *Anopheles* OR-positive cells has been examined (Schultze et al., 2013; Schymura et al., 2010), the population and the distribution patterns of IRs in the antenna remains poorly characterized. Here, we performed whole mount fluorescent in situ hybridization (WM-FISH) of the antenna for the majority of antennal IRs. We provide a detailed spatial antennal map of the location of IR-expressing neurons across the antennae, highlighting a global organization of IRs into proximal and distal regions of the antenna. This work presents the first, to our knowledge, comprehensive organization map of IR-expressing neurons in mosquitoes. The abundance, spatial localization, and co-expression pattern of cells expressing a defined IR could offer insights into their functional roles in mediating the detection of behaviorally relevant odors.

## Results

### Comparative transcriptomic profiles of *An. coluzzii* IRs in the main olfactory appendage

We performed transcriptomic analysis on the female *An. coluzzii* (strain N’Gousso) antennae, the main olfactory appendage (Fig 1A). RNAseq data for IRs were retrieved from transcriptomic data reported from prior studies (Athrey et al., 2017; Maguire et al., 2022; Pitts et al., 2017; Rinker et al., 2013b). A total of 44 IRs were predicted in the *An. gambiae* genome comprising 3 IR co-receptors and 41 tuning IRs (Fig 1B). We next compared antennal transcriptome RNAseq data across the 4 studies (Fig 1C). As expected, abundant expression levels for the IR-coreceptors (IR8a, IR25a, and IR76b) was consistently identified (orange labels in Fig 1C). A cut-off transcript level (>10 RPKM reported from at least two datasets) was set to define highly expressed IRs. Arrangement of IRs based on transcript abundance revealed the expression of 20 highly expressed IRs, whereas a total of 24 lowly expressed or undetected IRs (<10 RPKM from two or more datasets) were reported across all 4 antennal transcriptome datasets (Fig 1C). The data from Maguire et al., (2022) and Rinker et al., (2013) identified IR25a as the most abundant IR in the antenna. Other studies reported that IR76b transcript levels exceeded other IR co-receptors (Athrey et al., 2017; Pitts et al., 2017). Interestingly, the IR75l tuning receptor transcript level was more enriched than IR8a coreceptor in two independent studies (Pitts et al., 2011; Rinker et al., 2013b). Differences in transcript values might be explained by differences in tissue preparation, data acquisition or bioinformatic analysis. The IRs not abundant in antennal tissues might still be expressed in other tissues. The transcript enrichment of tuning IRs in the antenna, such as IR75l, IR41t.2, IR7w, and IR41c, might suggest an important functional role for these IRs in this main olfactory appendage.

**Fig 1:**
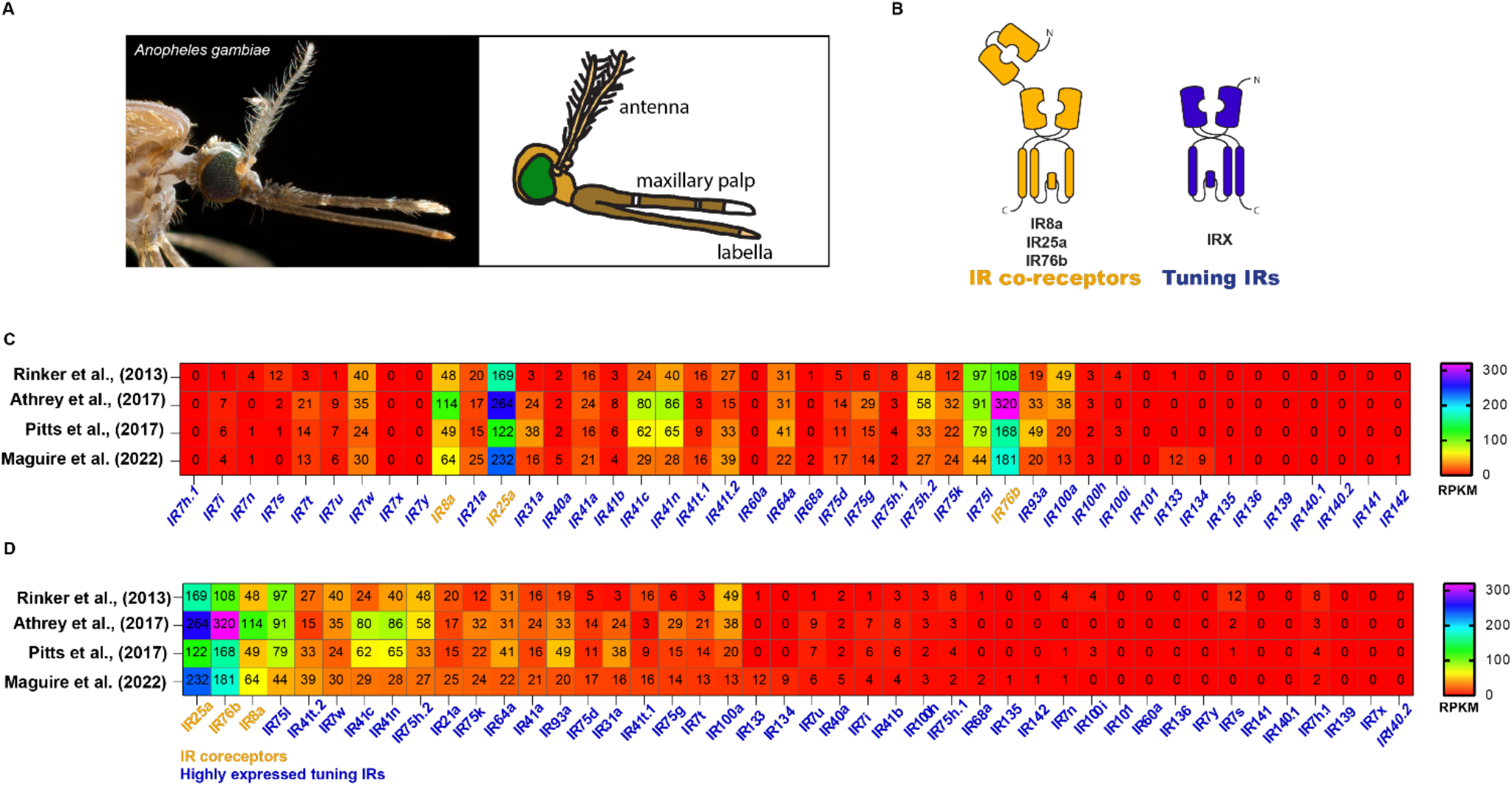
Transcript abundance of olfactory ionotropic receptors in the main olfactory appendage of *Anopheles gambiae* mosquitoes. **A**. A side view of a female *Anopheles gambiae* mosquito showing the olfactory appendages (antenna, maxillary palp, and labella) involved in odor detection and taste. Photo by Alex Wild. **B**. Schematic of *Anopheles* ionotropic receptors (IR)s showing the structures of IR co-receptors and tuning (odor-binding) IRs. IRs are ligand-gated ion channels capable of forming heteromeric functional complexes. Each tuning IRX requires at least one or more IR co-receptors (IR8a, IR25a and IR76b) to function. **C**. Heat map showing the transcript levels of all IRs predicted from the *Anopheles* genome. Data is compared across four independent studies. **D**. Heat map arranged by transcript abundance of IRs in the female antennae.

### Spatial expression of antennal IRs revealed by fluorescent in situ hybridization

RNAseq data reports the abundance of an IR in bulk antennal tissues but does not reveal the location or number of the IR-expressing neurons across the antenna. For example, highly enriched antennal IR transcripts might be due to one small antennal segment (flagellomere) containing many neurons expressing an IR, to many IR-expressing neurons spread across all 13 antennal flagellomeres, or to a few IR neurons expressing high levels of the IR. To map IR-expressing neurons, we performed whole-mount fluorescent in situ hybridization (WM-FISH) on antennal tissue using fluorescently labeled probes of target IRs. We first performed immunostaining on the 7 most abundant IRs across four independent transcriptomic data obtained from the female antennae (Fig 1D). The probes for 4 tuning IRs (Ir7w, Ir41c, Ir41t.2, Ir75l) and 3 co-receptor IRs (Ir8a, Ir76b, Ir25a) detected the localization of these IRs across the flagellomeres in the female and male antenna (female: Fig 2A-G; male: Fig 2-suppl 1). As expected, each flagellomere of the female antennae was densely populated with IR co-receptors (Fig 2E-G). Interestingly, spatial mapping of the abundant tuning IRs revealed differential expression across the antenna. Clusters of Ir7w, Ir41c, Ir41t.2 expressing neurons were found primarily in distal flagellomeres, while Ir75l was found only in 1-2 olfactory neurons in proximal flagellomeres (Fig 2A-D).

**Fig 2:**
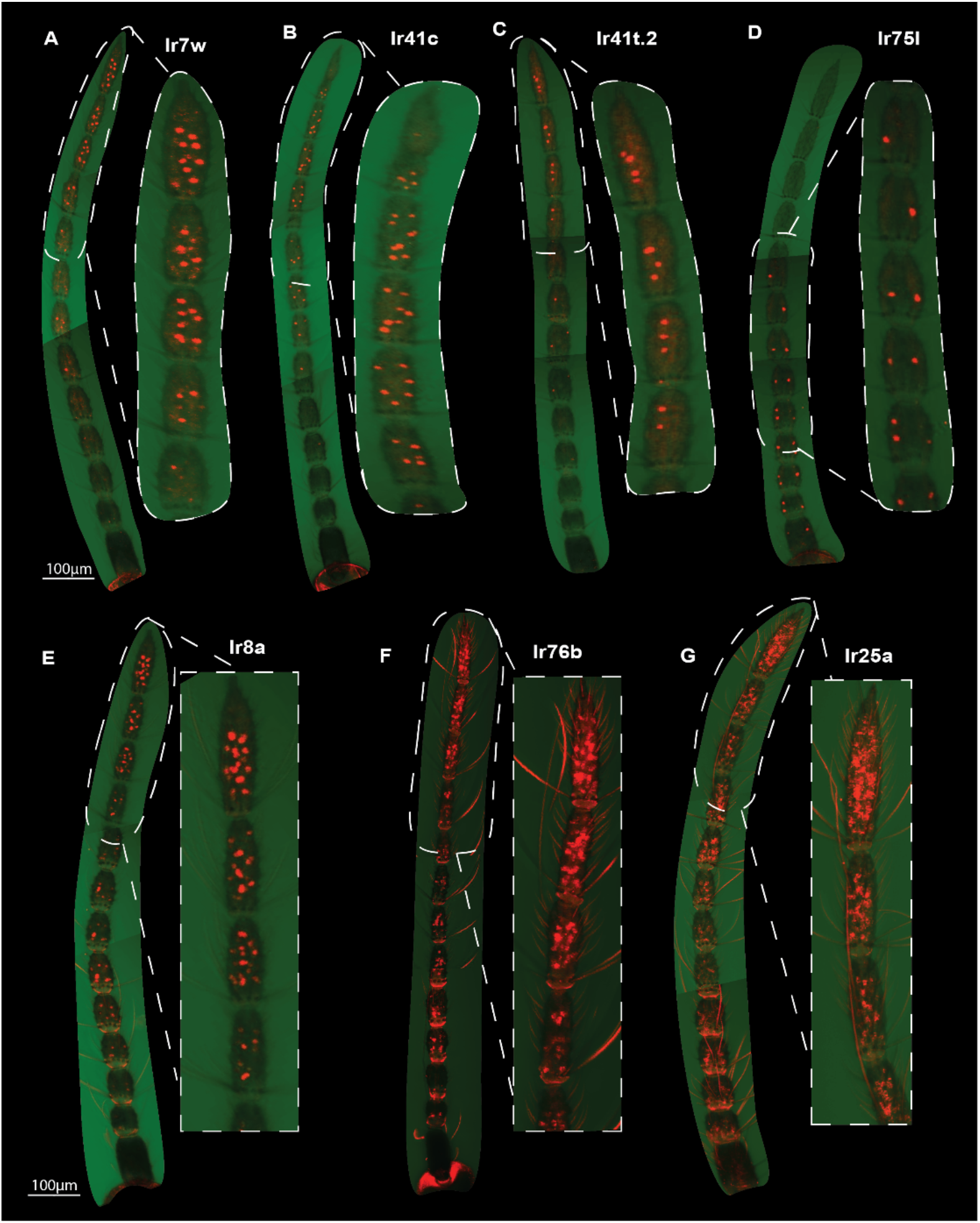
Topography of highly expressed IRs in the female *Anopheles coluzzii antenna*. Whole mount fluorescent in situ images of female antenna. Z-stack images from three different sections of the antenna were captured at 25X magnification and combined to form a single antennal image. Probes were generated for highly expressed tuning IRs: **A)** IR7w, **B)** IR41c, **C)** IR41t.2, **D)** IR75l, and IR coreceptors **E)** IR8a, **F)** IR76b, and **G)** IR25a. The antennal region demarcated by a white dashed line is also shown magnified.

The female antenna is morphologically different from the male antenna (Fig 3A-B). Although both are segmented into 13 flagellomeres, male antennae contain long (presumably mechanosensory) fibrillae bristles on flagellomeres 1-10 not present in the female. To categorize the topography of IRs in the female antenna, we grouped the flagellomeres into proximal (1-6) and distal (7-13) regions (Fig 3A). Statistical analysis revealed IR25a and IR76b (but not IR8a) as co-receptors more abundantly localized to the distal region (Fig 3C). This might reflect a higher density of IR-expressing neurons in distal flagellomeres than in proximal flagellomeres. Notably, IR75l-positive cells were robustly localized to the proximal region compared to the distal part (P=0.0003) (Fig 3 suppl 1). On the other hand, we identified more cells expressing IR7w, IR41t.2, and IR41c tuning IRs in the distal region (Fig 3 suppl 1).

**Fig 3:**
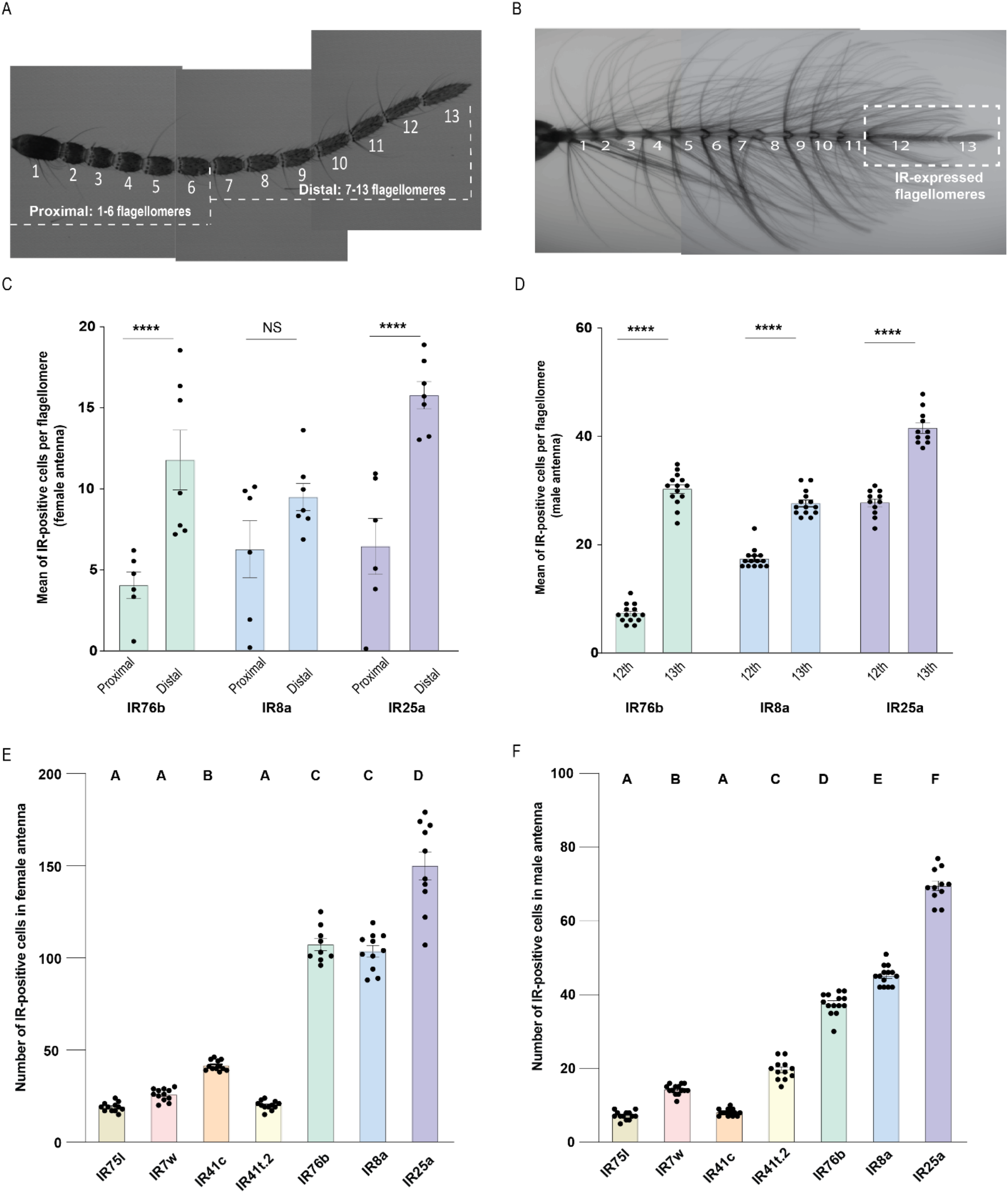
Distribution patterns of IRs across the male and female antennal flagellomeres. Confocal Z-stack images of *An. coluzzii* antenna. The numbers represent flagellomeres marked from the most proximal (1) to the most distal (13). **A)** Female antennal image showing the 13 flagellomeres grouped into proximal and distal regions. **B)** Male antennal image with the flagellomeres containing IR-expressing neurons marked with a dashed box. **C)** Bar graph showing the mean number of IR co-receptors per antennal region. Data was analyzed by two-sample T-test to compare average number of IR-positive cells detected in the proximal and distal region (NS= not significant; ****=P<0.00001; n= 6-7). **D)** Average number of cells detected in the 12^th^ and 13^th^ flagellomere from male antenna. Data was analyzed by two-sample T-test to compare average number of IR-positive cells detected in the proximal and distal region (****=P<0.00001; n= 11-14). **E)** Number of cells expressing the 4 most highly expressed tuning IRs and the 3 IR co-receptors in the female antenna. IRs with different letters are significantly different when analyzed by one-way ANOVA (P<0.0001; n= 9-12). **F)** Number of cells expressing the 4 most highly expressed tuning IRs and the 3 IR co-receptors in the male antenna. IRs with different letters are significantly different when analyzed by one-way ANOVA (P<0.0001; n= 11-14).

Unlike the female antenna, IR-positive cells were all physically localized to the 12^th^ and 13^th^ flagellomeres and undetectable in proximal segments of the male antenna. The 13^th^ flagellomere of the male antenna was more enriched with IR co-receptors than the 12^th^ segment (Fig 3D). Consistent with the robust transcript abundance of IR co-receptors (Fig 1D), the number of cells expressing IR co-receptors exceeded the tuning IRs in the female (Fig 3E) and male (Fig 3F) antennae. Among the highly expressed tuning IRs examined in the female antennae, IR41c-positive cells were more robustly detected compared to other tuning IRs, whereas IR41t.2 expressing cells were the most enriched tuning IR in the male antennae (Fig 3E-F). The abundance of IR41c-expressing cells in the female antennae could indicate their importance in detecting a crucial host odor cue. The male enriched IR41t.2 might similarly be important for male behaviors like nectar seeking or mate finding. A comparison between the number of IR-expressing cells and transcriptomics data suggested that some IRs might be highly expressed in a small number of neurons (Table 1). For example, IR75l transcript levels were reported to be more abundant than IR8a (Pitts et al., 2017; Rinker et al., 2013b), but the number of cells expressing IR75l was fewer compared to IR8a in both male and female antennae (Fig 1D, Fig 2, Fig 2 Supp 1). This suggests IR75l might be highly expressed in a small set of neurons, while the more densely populated IR8a-positive cells express relatively lower levels of the IR8a transcript. IR93a transcripts per olfactory neuron similarly appeared high based on transcript levels and the number of IR93a-expressing neurons (Table 1).

**Table 1.**
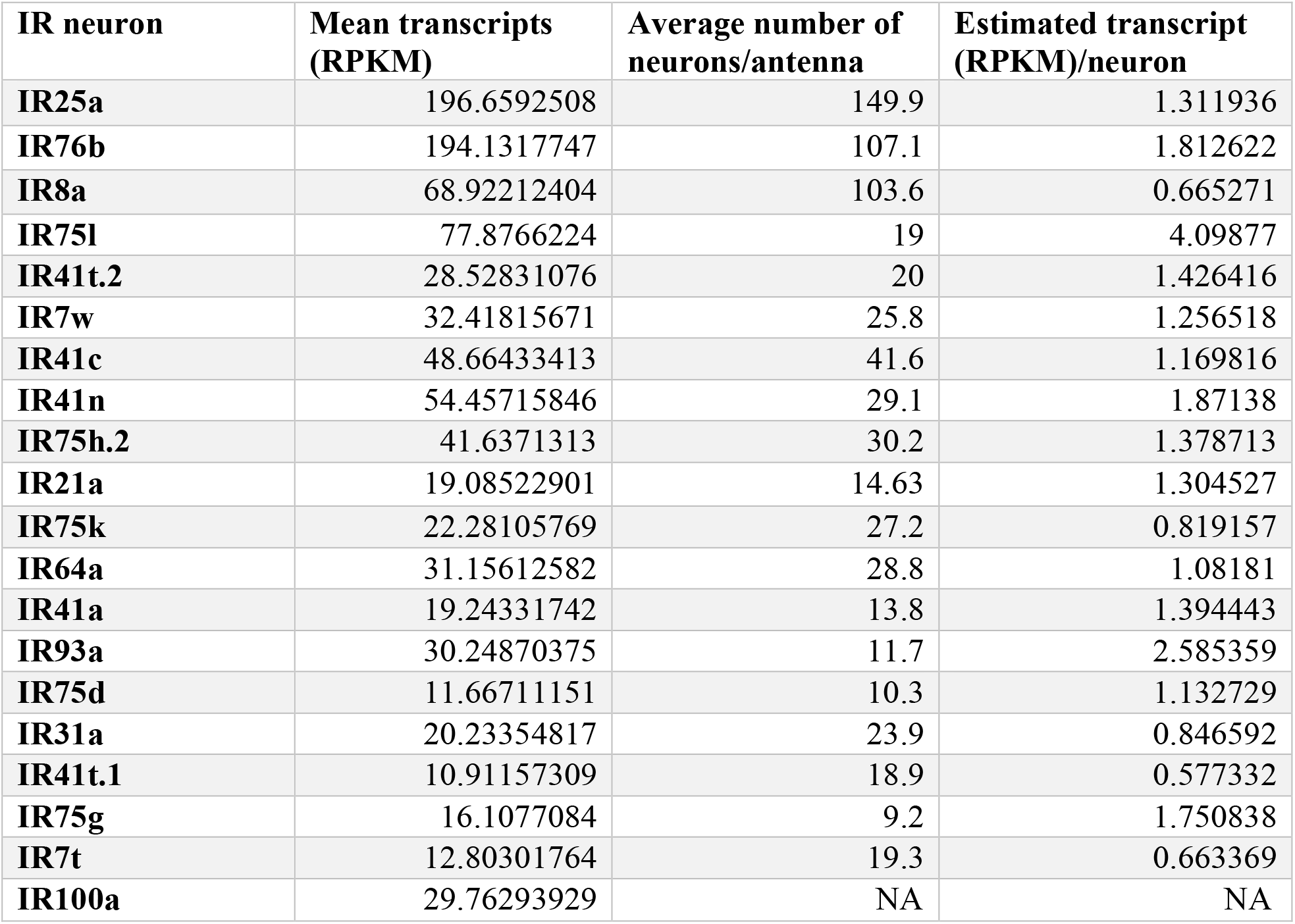
Estimate of IR transcript abundance per IR-expressing neuron in the female antenna. **Related to Figure 1 and 3**. Mean transcripts were averaged over 4 transcriptome datasets (Athrey et al., 2017; Maguire et al., 2022; Pitts et al., 2017; Rinker et al., 2013b) while the average number of neurons in an antenna was retrieved from the fluorescent in situ data. The IR expression level per antennal neuron was calculated by dividing the mean transcripts (Reads Per Kilobase of transcript, per Million mapped reads) by the average number of IRX-expressing neurons. IR100a was not detected by in situ, hence noted as NA (not applicable).

| IR neuron | Mean transcripts (RPKM) | Average number of neurons/antenna | Estimated transcript (RPKM)/neuron |
| --- | --- | --- | --- |
| IR25a | 196.6592508 | 149.9 | 1.311936 |
| IR76b | 194.1317747 | 107.1 | 1.812622 |
| IR8a | 68.92212404 | 103.6 | 0.665271 |
| IR75l | 77.8766224 | 19 | 4.09877 |
| IR41t.2 | 28.52831076 | 20 | 1.426416 |
| IR7w | 32.41815671 | 25.8 | 1.256518 |
| IR41c | 48.66433413 | 41.6 | 1.169816 |
| IR41n | 54.45715846 | 29.1 | 1.87138 |
| IR75h.2 | 41.6371313 | 30.2 | 1.378713 |
| IR21a | 19.08522901 | 14.63 | 1.304527 |
| IR75k | 22.28105769 | 27.2 | 0.819157 |
| IR64a | 31.15612582 | 28.8 | 1.08181 |
| IR41a | 19.24331742 | 13.8 | 1.394443 |
| IR93a | 30.24870375 | 11.7 | 2.585359 |
| IR75d | 11.66711151 | 10.3 | 1.132729 |
| IR31a | 20.23354817 | 23.9 | 0.846592 |
| IR41t.1 | 10.91157309 | 18.9 | 0.577332 |
| IR75g | 16.1077084 | 9.2 | 1.750838 |
| IR7t | 12.80301764 | 19.3 | 0.663369 |
| IR100a | 29.76293929 | NA | NA |

### *Anophele*s IRs are expressed in stereotyped topographical regions across the antenna

We next generated 20 additional in situ probes and performed whole mount fluorescent in situs to map the spatial distribution for all antennal-expressed IRs as reported by transcriptomic data. In addition to the proximally localized IR75l-positive cells (Fig 3 supp 1), we detected additional IRs (IR41a and IR75d) that were enriched in the proximal region (Fig 4A-B). The majority of antennal IR-expressing cells were consistently abundant in the 12^th^ and 13^th^ flagellomeres of the female antenna (Fig 4A-B). Notably, approximately 80% of IR93a-expressing cells detected by in situ were localized to the 13^th^ flagellomere (Fig 4B). In a few samples, we detected the expression of IR75l and IR76b in the 1st flagellomere (Fig 4C).

**Fig 4:**
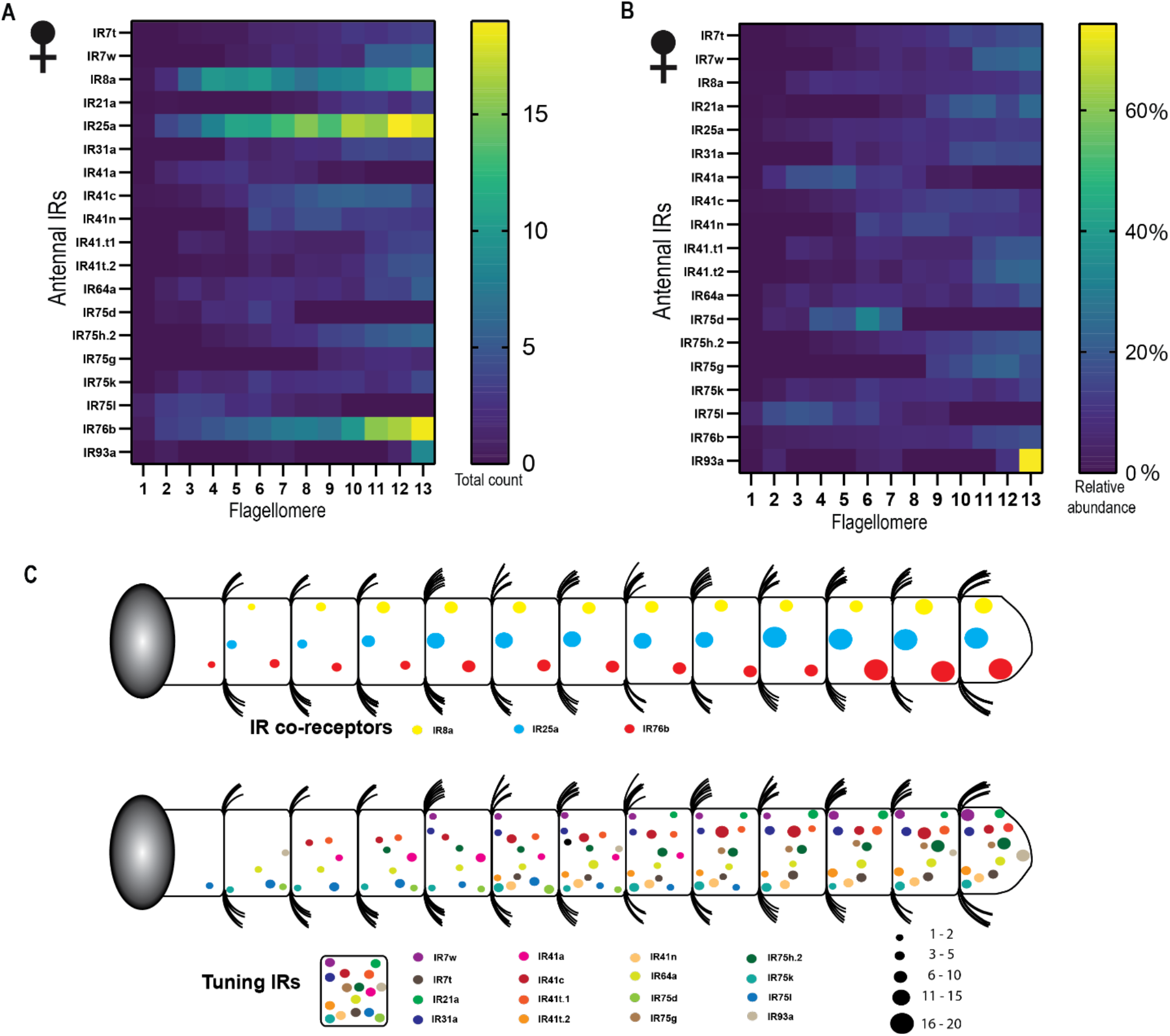
A spatial map of IR expression and distribution in the female antenna. **A)** Heat map representing the number of IR-expressing cells per flagellomere. **B)** Relative abundance of IR-positive cells across flagellomeres. **C)** Cartoon model showing the spatial map of IRs across the 13 flagellomeres. The size of the circle corresponds to the number of cells identified per flagellomere. The three IR co-receptors are shown separately from the tuning IRs. Each color represents a different IR. The key serves as a reference guide to help identify the tuning IRs found in a flagellomere; it does not indicate their relative spatial position in a flagellomere.

While the number of IRX-positive neurons per flagellomere was consistent from animal to animal, the position of IRX-positive cells within a flagellomere appeared to be more variable. Although RNAseq data detected IR100a in the female antenna, we could not detect cells expressing IR100a by whole mount fluorescence in situs. It is possible that IR100a expression level in a neuron was below the detection limit of our in situ protocol. Given this observation, we did not generate probes for IRs with transcript levels below IR100a.

We also analyzed the abundance of a limited set of IRs across the male antenna (Fig 4 Supp Fig 1). Consistent with the unique expression pattern of IR75l in the proximal region of the female antennae, the male antenna contained IR75l-positive cells only in the 12^th^ antennal segment, while the 13^th^ segment was devoid of IR75l cells (Fig 4 Supp fig 1). In general, more IR-expressing neurons were localized to the distal 13^th^ flagellomere than in the 12^th^ flagellomere (Fig 4 Supp Fig 1). It should be noted that the abundant IRs examined here by in situs were based on female antennal RNAseq expression studies; male specific antennal RNAseq might identify a different set of abundant IRs that warrant further study.

### Double immunofluorescence reveals the co-receptors for highly expressed tuning IRs

Tuning IRs form a functional complex with at least 1 or more co-receptors (Abuin et al., 2011; Ai et al., 2013), which might influence their functional properties (Task et al., 2022). To examine this, we utilized two probe whole mount fluorescent in situs to identify which IR co-receptors were co-expressed in the same cells as the 4 highly expressed tuning IRs. We identified the co-expression patterns of IR7w, IR41c, IR41t.2, and IR75l with IR8a, IR25a, and IR76b (Fig 5A-C). The IR41 clade of IRXs includes IR41c and IR41t.2 which showed similar co-expression patterns and did not co-label with IR8a (Fig 5A). We observed partial overlap between IR7w-positive cells and IR8a-expressing cells (Fig 5A). In contrast, IR75l expressing cells completely overlapped with IR8a-postive cells (Fig 5D). Co-localization of *An. coluzzii* IR7w and IR75l suggests they may be tuned to acids, given *Aedes* IR8a (Raji et al., 2019) and *Drosophila* IR8a expressing neurons (Ai et al., 2013) have been reported to detect acids. Notably, all the 4 tuning IRs examined in the study completely overlapped with IR25a-positive cells (Fig 5B & D). This observation is consistent with the premise that IR25a may express in most (if not all) IRX-expressing neurons. Recent studies have shown that IR76b neurons are responsive to amines (Vulpe and Menuz, 2021; Ye et al., 2021). Likewise, *Drosophila* IR25a synergistically interact with IR76b to elicit amine-evoked responses (Benton et al., 2009). The distinct co-expression patterns of IR41c and IR41t.2 with IR25a and IR76b, but not IR8a (Fig 5A-D), could suggest a role for these tuning IRs in the detection of amine chemical compounds.

**Fig 5:**
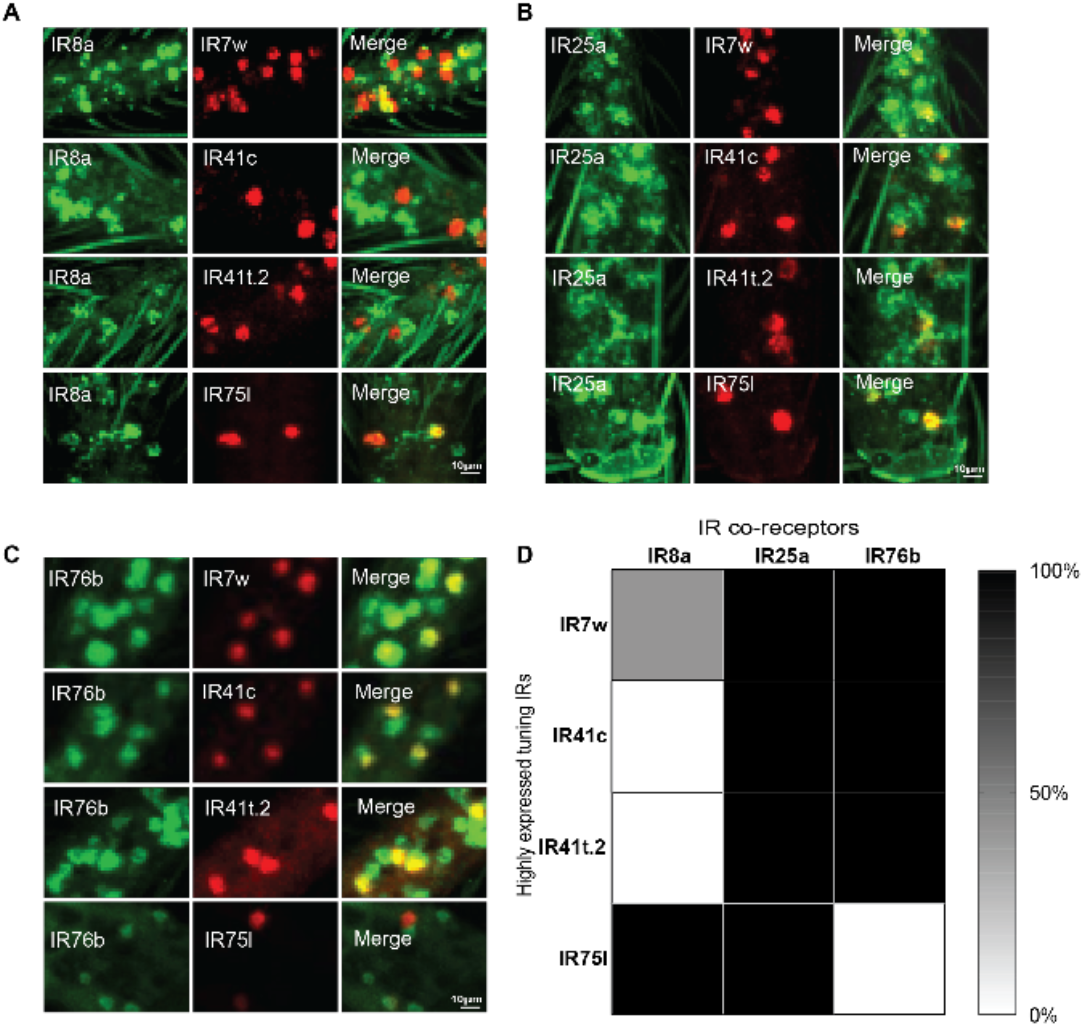
Co-expression patterns of highly expressed tuning IRs with IR co-receptors. In situ images showing co-localization of the 4 highly expressed tuning IRs with **A)** IR8a, **B)** IR25a, and **C)** IR76b co-receptors. Tuning IR probes were conjugated to the Alexa 647 fluorophore (red) while IR co-receptor probes were linked to the Alexa 488 fluorophore (green). **D)** Summary heat map showing the co-expression patterns of the 4 tuning IRs with IR co-receptors. The percent co-expression was calculated as the percent of tuning IR-positive cells that co-labeled with the IR co-receptor(s), n=7-9.

## Discussion

We present here a comprehensive spatial map of IR-expressing neurons across the entire *Anopheles* antenna. IRs were most abundant in the distal flagellomere in both female (Fig 4c) and male mosquitoes (Fig 4 suppl. Fig 1c), suggesting distal flagellomere to be the most sensitive region for detecting volatile cues. Nonetheless, we also identified IRXs whose expressing neurons were primarily localized only to proximal (IR41a, IR75d, IR75l) or distal (IR7w, IR21a, IR41n, IR41.t2, IR75g, IR93a) antennal flagellomeres. These data suggest an additional level of organization may exist separating the antenna into proximal and distal domains. If these domains similarly reflect discrete olfactory signaling domains remains to be explored.

Of the 44 IRs identified in the *Anopheles* genome, nearly half had elevated transcript levels in the antenna. Bulk antennal RNAseq data suggests weak transcript expression for some of these tuning IRs; however, we could not detect neurons expressing these IRs by in situ hybridization. This might reflect differences in the sensitivity of these two approaches. RNAseq, as a bulk measure, might be able to identify IRX transcripts that are weakly expressed across many neurons, whereas such low expression of IRX transcripts per neuron might be below the level of detection of our in situ method. Alternatively, RNAseq might be able to identify IRs expressed in small neuronal populations resistant to the in situ protocol.

### The flagellomere as an organizing antennal unit of IRs in the *Anopheles* antennae

By mapping the majority of tuning IRXs to olfactory neurons across many antennae, we can reveal if patterns of olfactory neuron organization exist within a flagellomere. We observed that the number of IRX-expressing cells appeared stereotyped from antenna to antenna; that is, the same number of IRX-expressing neurons were consistently found across many samples. Interestingly, the exact position of an IRX-expressing neuron within a flagellomere was variable. This suggests the flagellomere is an organizing unit for olfactory neurons, but olfactory neurons do not occupy fixed locations within each flagellomere.

A neuron might be able to increase its sensitivity to sensory cues by increasing expression of its IRX receptor. While it remains to be determined if transcript levels directly relate to protein abundance, our work identified a couple of IR-expressing neurons that appear to be highly expressed. In particular, IR75l and IR93a appear to be at least two-fold more highly expressed in their respective neurons compared to other IRs. IR93a has been linked to temperature and humidity-sensing in *Drosophila* and *Anopheles* (Knecht et al., 2016; Laursen et al., 2022), and an increased abundance might be required to mediate this affect. The functional role for IR75l remains to be determined. It also remains possible that additional antennal neurons express these IRX receptors at levels below the detection of our in situ analyses.

Transcriptome analyses suggest that mosquito olfactory receptor expression might change with the age of the animal (Omondi et al., 2019; Tallon et al., 2019). Recent work examining *Anopheles coluzzii* ORs suggest that changes in OR expression between 1-8 days after pupal eclosion might reflect changes to the number of neurons expressing a particular OR (Maguire et al., 2022). Such changes in olfactory receptor expression might be a mechanism to guide olfactory behaviors. If these expression studies on ORs extends to IRs remains to be determined. We have presented a snapshot of the IR spatial map for the *Anopheles* antenna during day 5-7 post eclosion which would coincide with host-seeking behaviors in the *An. coluzzii* species (Omondi et al., 2019). It would be interesting to determine how the snapshot of IR-expressing neurons at this stage might differ during other behavioral stages such as after blood feeding and during oviposition. It is possible that some IRs might even be absent during the adult stage we examined, and only expressed during discrete behaviors.

### Combinatorial co-expression patterns of *Anopheles* IRs offer functional insights

The function of a tuning IRX towards amines or acids can often be predicted by which IR co-receptor it complexes with. Functional analysis showed that IR64a is an acid tuning receptor that forms a functional complex with IR8a to elicit acid-evoked responses (Ai et al., 2013). Immunoprecipitation and immunostaining of *Drosophila* IR64a and the IR8a co-receptor protein further revealed that both receptors physically interact and co-express (Ai et al., 2013). In *Ae aegypti*, IR8a is required for detecting carboxylic acids (Raji et al., 2019) - a role conserved in *Drosophila* (Ai et al., 2013). *Drosophila* IR76b functions alongside IR25a as co-receptors necessary for amine detection (Vulpe and Menuz, 2021). Similarly, *An. coluzzii* IR76b functions as a coreceptor in neurons responsive to amines (Ye et al., 2021). In *Anopheles* larvae, RNAi mediated silencing of IR76b caused impaired response to butylamine (Liu et al., 2010). These data suggest that IRX co-expression with IR8a likely suggests responses towards acids, whereas co-expression with IR76b suggests a functional response towards amines.

By in situs analyses, we found IR41c-expressing neurons to co-localize with IR25a and IR76b, but not IR8a, suggesting IR41c might function to respond to amines. Indeed, heterologous expression of Ir41c (along with co-receptors Ir76b and Ir25a) in *Xenopus* oocyctes revealed Ir41c might mediate responses to the amines 3-methylpiperidine, 3-pyrroline, and pyrrolidine (Pitts et al., 2017). The exclusive co-expression of IR75l with the IR8a coreceptor suggests it will play a role in detecting acids. IR75l is a closely related member of *Anopheles* IR75k in the IR75 clade. The response profile of IR75k was determined by heterologous expression of IR8a and IR75k in *Xenopus* oocytes (Pitts et al., 2017). This study revealed that IR75K could detect a range of six to ten chain length carboxylic acids, with peak responses to octanoic (8 carbon) and nonanoic (9 carbon) carboxylic acids. This suggests that IR75l-expressing neurons might also respond to carboxylic acids.

The expression of the 3 IR co-receptors with a tuning IR might be utilized to alter the signal sensitivity of a neuron expressing that particular combination of the IRX-IRCO complex. For example, when *Drosophila* IR76a was co-expressed with IR25a and IR76b in an empty neuron system, this combination evoked robust responses to phenylethyl amine which was reduced by ∼50% when co-expressed with either IR25a or IR76b alone (Abuin et al., 2011). These results suggest functional insights might be gained by similar in situ studies presented here in which the co-receptor expression patterns for all tuning IRs across a mosquito antenna are cataloged.

## ACKNOWLEDGEMENTS

We thank all the members of Potter lab for helpful discussions. This work was supported by grants from the National Institutes of Health to C.J.P. (NIAID R01Al137078), the Department of Defense to C.J.P. (W81XWH-17-PRMRP), a Johns Hopkins Postdoctoral Research Accelerator Award to J.I.R, and a Johns Hopkins Malaria Research Institute Postdoctoral Fellowship to J.I.R. We thank the Johns Hopkins Malaria Research Institute and Bloomberg Philanthropies for their support.

## Author Contributions

**Conceptualization**: J.I.R. and C.J.P.; **Methodology**: J.I.R. and C.J.P.; **Formal Analysis**: J.I.R. **Investigation**: J.I.R.; **Writing, Original Draft**: J.I.R. and C.J.P.; **Writing, Review & Editing**: J.I.R. and C.J.P.; **Visualization**: J.I.R. and C.J.P.; **Supervision**: C.J.P.; **Funding Acquisition**: C.J.P.

## Materials and Methods Method details

### Mosquito rearing

*Anopheles coluzzii* (Ngousso strain; formerly *Anopheles gambiae* M form), were maintained under 12 h light/dark cycle at 28°C, and 72±2% relative humidity. The larvae were fed ad libitum on TetraMin Tropical Flakes and Purina Cat pellets. The emerged adult mosquitoes were raised with 10% sucrose solution. To generate eggs, adult female mosquitoes were fed on anaesthetized mice or defibrinated sheep blood.

### Whole-mount fluorescent in situ hybridization (WM-FISH)

This assay was performed as previously described with some modifications (Younger et al., 2020; Maguire et al., 2022). All in situ probe reagents used were purchased from Molecular Instruments, Inc. Mosquito antennae aged 5-7 days post eclosion were dissected into CCD buffer (50 units of chitinase, 1000 units of chymotrypsin (25 mg of 40 units/mg), 10 mL HEPES larval buffer (119 mM NaCl, 48 mM KCl, 2 mM CaCl_2_, 2 mM MgCl2, 25 mM HEPES), and 100 µl DMSO) on ice, then incubated for 25 minutes at 37°C on a rotator. Tissues were then pre-fixed in 4% paraformaldehyde (PFA) in PBT (1XPBS, 0.03% Triton X-100) for 24h at 4°C. Tissue was washed with 0.1% PBS-Tween on ice, incubated for 1 hour at RT in 80% methanol/20% DMSO. Tissues were incubated overnight at -20°C in absolute methanol. Using different series of graded Methanol/PBS-Tween (75% methanol, 50% methanol; 25% methanol; and 100% PBS-Tween in that order), tissues were rehydrated for 10 mins on ice and washed with PBS-Tween at room temperature. This was followed by incubation in Proteinase K solution (20ug/ml) for 30 mins and washed in PBS-Tween at room temperature. Tissues were post-fixed in 4% PFA in PBS-Tween at RT for 20 minutes. Tissue was washed in PBS Tween at room temperature three times for 15 mins per wash, then pre-hybridized in pre-heated probe hybridization buffer at 37°C for 30 minutes. Tissue was incubated in probe solution (8 pmol in hybridization buffer) at 37°C for two nights. Thereafter, tissue was washed at least 5 times in pre-heated probe wash buffer at 37°C, then washed at room temp in saline-sodium citrate dissolved in 1% Tween. Pre-amplification was performed using the amplification buffer at room temperature for 10 minutes, then incubated in the hairpin mixture (18 pmol snap-cooled hairpins in amplification buffer). Tissue was incubated overnight in the dark at room temperature. This was followed by a series of washes in saline-sodium citrate solution performed at room temperature to remove excess hairpins. After, tissue was gently dipped in droplets of SlowFade Diamond (ThermoFisher S36972) at least three times to clear excess wash solution before mounting in SlowFade Diamond.

### Data acquisition

Following in situs, slides containing whole antenna tissue were mounted and the IR-positive cells were counted manually under a Zeiss LSM 700 confocal microscope. Images were acquired at 512 × 512-pixel resolution with 0.58, 2.37 or 6.54 µm z-steps by using the Fluar 10x air M27, LCI 25x water immersion with Korr DIC M27. For illustration purposes, confocal images were exported as maximum intensity projection and processed in ImageJ and annotated on Adobe illustrator. Images were adjusted for brightness or contrast and unmodified.

### Statistical analysis

Data quantification and analysis was performed using the GraphPad Prism 9 software package (GraphPad Software, San Diego, CA). We performed two-sample T-tests and one-way ANOVA statistical tests in this study. Full statistical details can be found in the figure legends including statistical tests, number of samples and trials, and how statistical significance was determined.

### Key Resources Table

| Key Resources Table |  |  |  |  |
| --- | --- | --- | --- | --- |
| Reagent type (species) or resource | Designation | Source or reference | Identifiers | Additional information |
| commercial assay or kit | Probe sets | Molecular Instruments, Inc | HCR v3.0 | https://www.molecularinstruments.com/ |
| Software and Algorithms | GraphPad software | www.graphpad.com | GraphPad Prism 8.0 & 9.0 |  |
| Software and Algorithms | Fiji (ImageJ) | https://imagej.net/Fiji |  | Free image editing tool |
| Software and Algorithms | Adobe Illustrator | https://www.adobe.com/products/illustrator.html | Adobe illustrator CS6 | Adobe, Inc. |
| IR7w in situ probe | AGAP01346 (2002 nt) | Molecular Instruments, Inc | Probe Set ID: PRH968 | RNA in situ probes designed by Molecular Instruments based on coding sequence |
| IR41C in situ probe | AGAP012951 (2229 nt) | Molecular Instruments, Inc | Probe Set ID: PRH969 | RNA in situ probes designed by Molecular Instruments based on coding sequence |
| IR41t.2 in situ probe | AGAP012969 (2264 nt) | Molecular Instruments, Inc | Probe Set ID: PRH970 | RNA in situ probes designed by Molecular Instruments based on coding sequence |
| IR75l in situ probe | AGAP005466 (2349 nt) | Molecular Instruments, Inc | Probe Set ID: PRH971 | RNA in situ probes designed by Molecular Instruments based on coding sequence |
| IR25a in situ probe | AGAP010272 (4005 nt) | Molecular Instruments, Inc | Probe Set ID: PRK149 | RNA in situ probes designed by Molecular Instruments based on coding sequence |
| IR8a in situ probe | AGAP010411 (2568 nt) | Molecular Instruments, Inc | Probe Set ID: PRK150 | RNA in situ probes designed by Molecular Instruments based on coding sequence |
| IR76b in situ probe | AGAP011968 (7156 nt) | Molecular Instruments, Inc | Probe Set ID: PRI998 | RNA in situ probes designed by Molecular Instruments based on coding sequence |
| IR41n in situ probe | AGAP003531 (2282 nt) | Molecular Instruments, Inc | Probe Set ID: PRK696 | RNA in situ probes designed by Molecular Instruments based on coding sequence |
| IR75h.2 in situ probe | AGAP001812 (2291 nt) | Molecular Instruments, Inc | Probe Set ID: PRK697 | RNA in situ probes designed by Molecular Instruments based on coding sequence |
| IR21a in situ probe | AGAP008511 (3209 nt) | Molecular Instruments, Inc | Probe Set ID: PRK698 | RNA in situ probes designed by Molecular Instruments based on coding sequence |
| IR75k in situ probe | AGAP007498 (2172 nt) | Molecular Instruments, Inc | Probe Set ID: PRK699 | RNA in situ probes designed by Molecular Instruments based on coding sequence |
| IR64a in situ probe | AGAP004923 (3767 nt) | Molecular Instruments, Inc | Probe Set ID: PRK700 | RNA in situ probes designed by Molecular Instruments based on coding sequence |
| IR41a in situ probe | AGAP002904 (2872 nt) | Molecular Instruments, Inc | Probe Set ID: PRK974 | RNA in situ probes designed by Molecular Instruments based on coding sequence |
| IR93a in situ probe | AGAP000256 (5100 nt) | Molecular Instruments, Inc | Probe Set ID: PRK975 | RNA in situ probes designed by Molecular Instruments based on coding sequence |
| IR75d in situ probe | AGAP004969 (1824 nt) | Molecular Instruments, Inc | Probe Set ID: PRK976 | RNA in situ probes designed by Molecular Instruments based on coding sequence |
| IR31a in situ probe | AGAP009014 (2660 nt) | Molecular Instruments, Inc | Probe Set ID: PRK977 | RNA in situ probes designed by Molecular Instruments based on coding sequence |
| IR41t.1 in situ probe | AGAP004432 (2333 nt) | Molecular Instruments, Inc | Probe Set ID: PRK978 | RNA in situ probes designed by Molecular Instruments based on coding sequence |
| IR75g in situ probe | AGAP013085 (3263 nt) | Molecular Instruments, Inc | Probe Set ID: PRL354 | RNA in situ probes designed by Molecular |
|  |  |  |  | Instruments based on coding sequence |
| IR7t in situ probe | AGAP002763 (1978 nt) | Molecular Instruments, Inc | Probe Set ID: PRL355 | RNA in situ probes designed by Molecular Instruments based on coding sequence |
| IR100a in situ probe | AGAP000140 (3606 nt) | Molecular Instruments, Inc | Probe Set ID: PRL356 | RNA in situ probes designed by Molecular Instruments based on coding sequence |

## Figures and Tables

**Fig 2 Suppl 1:**
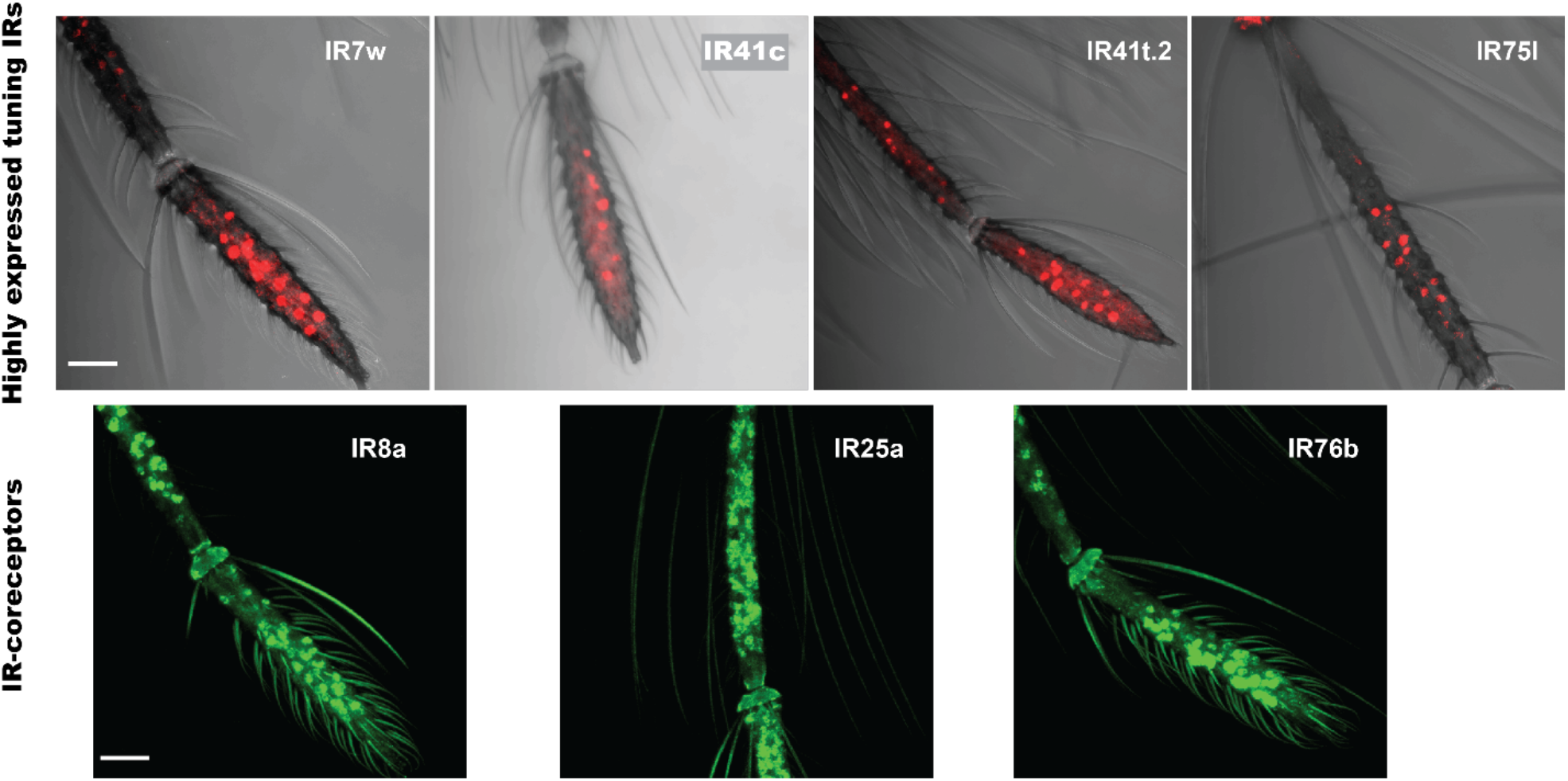
Topography of highly expressed IRs in the male *Anopheles coluzzii antenna*. Confocal images showing full z-stacks of a male *An. coluzzii* antenna. Whole-mount immunostaining was performed for highly expressed tuning IRs and IR co-receptors. For tuning IRs, probes were conjugated to Alexa 647 while IR co-receptor probes were conjugated to Alexa 488. Scale bars represent 50µm.

**Fig 3 suppl 1:**
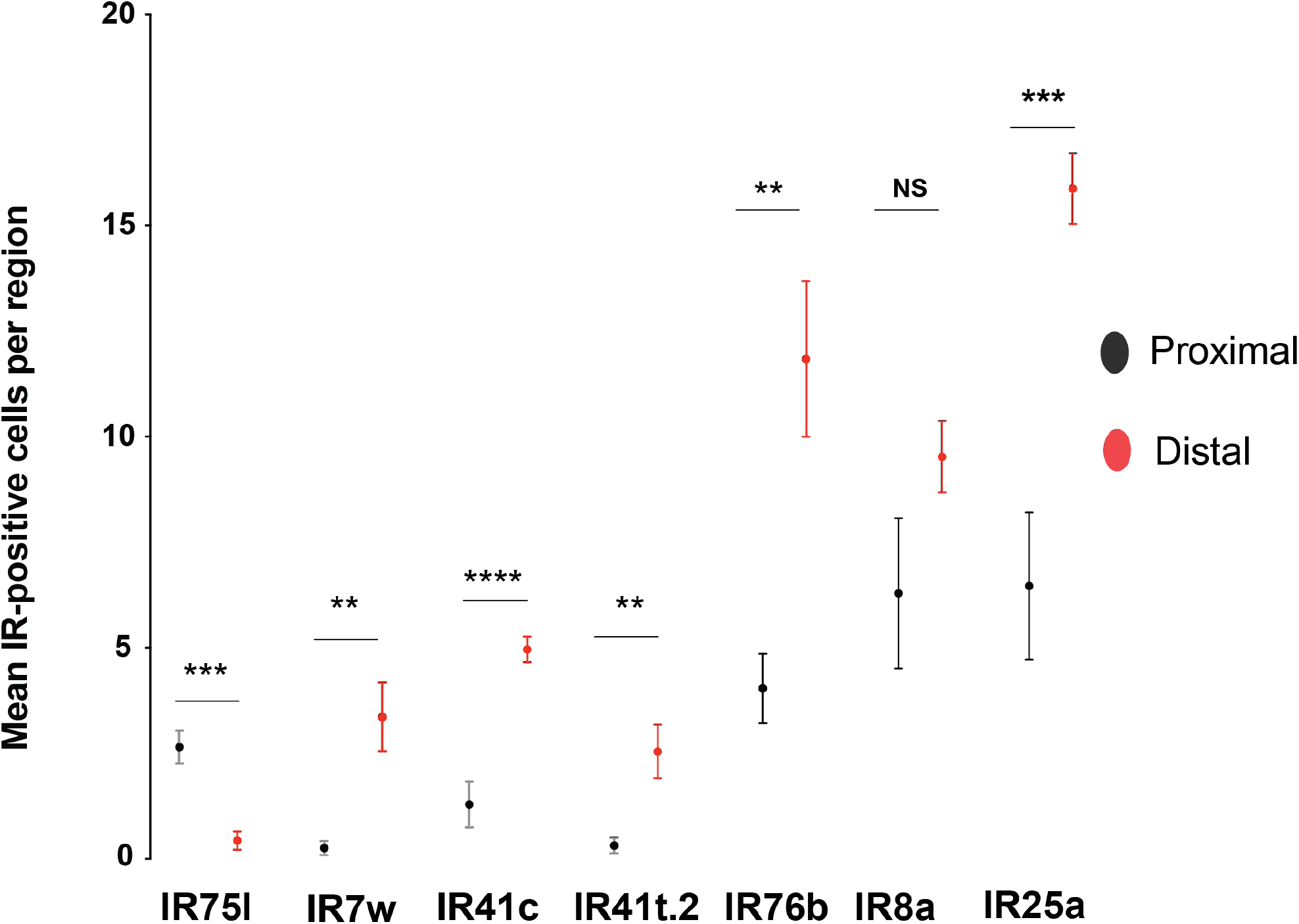
IR expression in proximal and distal regions of the female antenna. The mean number of tuning IRs (IR75l, IR7w, IR41c, IR41t.2) and IR co-receptors (IR76b, IR8a, IR25a) per antennal region is shown. Data was analyzed by T-test to compare average number of IR-positive cells detected in the proximal and distal region (NS= not significant; **= P<0.001; ***=P<0.0001; ****=P<0.00001; n= 6-7).

**Fig 4 Suppl 1:**
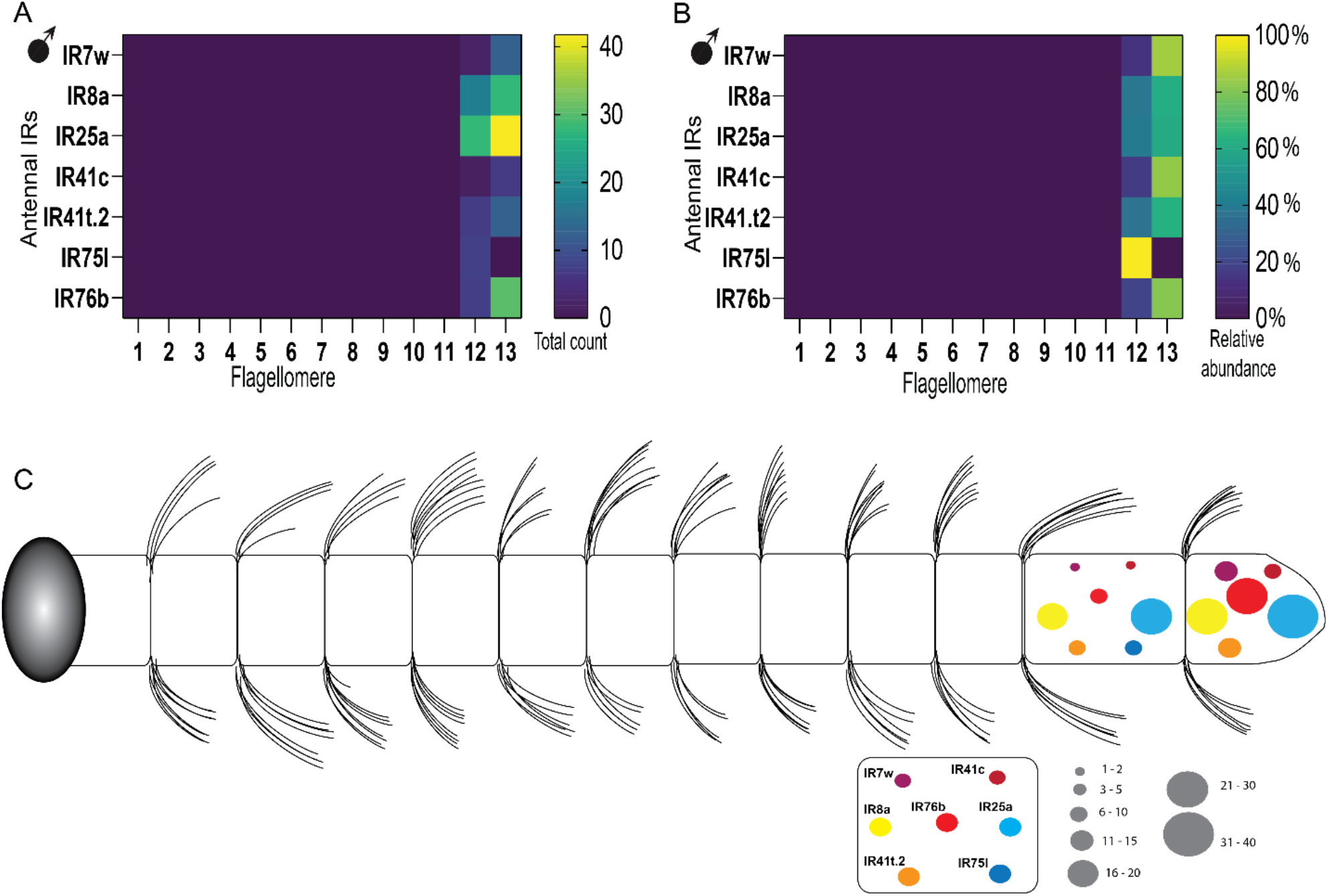
A spatial map of IR expression and relative distribution in the male antenna. **A)** Heat map representing the number of IR-positive cells per flagellomere (n=9-14). **B)** Heat map showing the relative abundance of IRs in a flagellomere compared across an antenna (n=9-14). C) A cartoon model showing a spatial map of expressed IRs in the male antenna of *An. coluzzii*.

**Fig 5 suppl 1:**
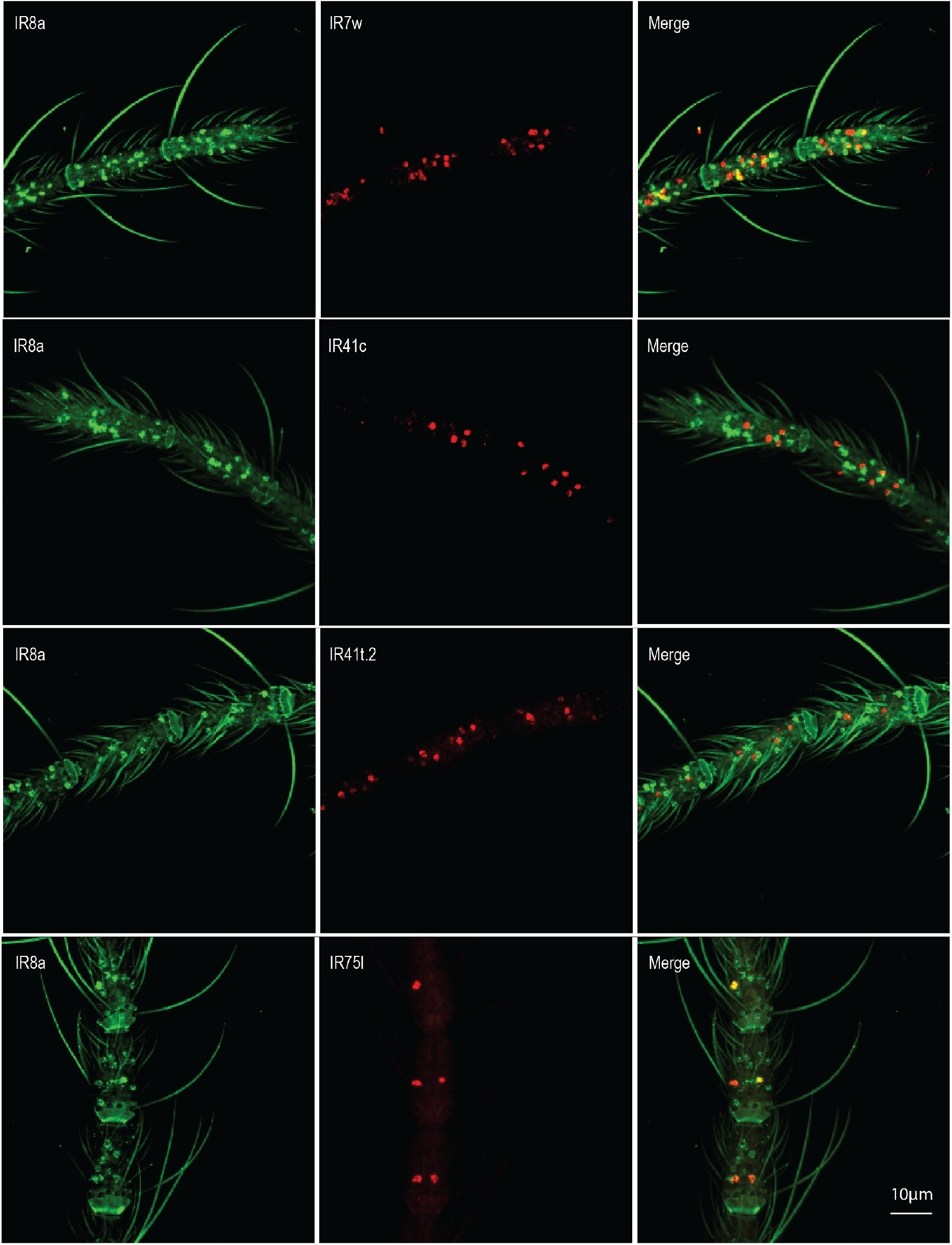
Co-expression of highly expressed tuning IRs with IR8a. In situ images showing co-expression of highly expressed tuning IRs with IR8a across the flagellomeres. Tuning IR probes were conjugated to the Alexa 647 fluorophore (red) while the IR8a co-receptor probe was linked to the Alexa 488 fluorophore (green).

**Fig 5 suppl 2:**
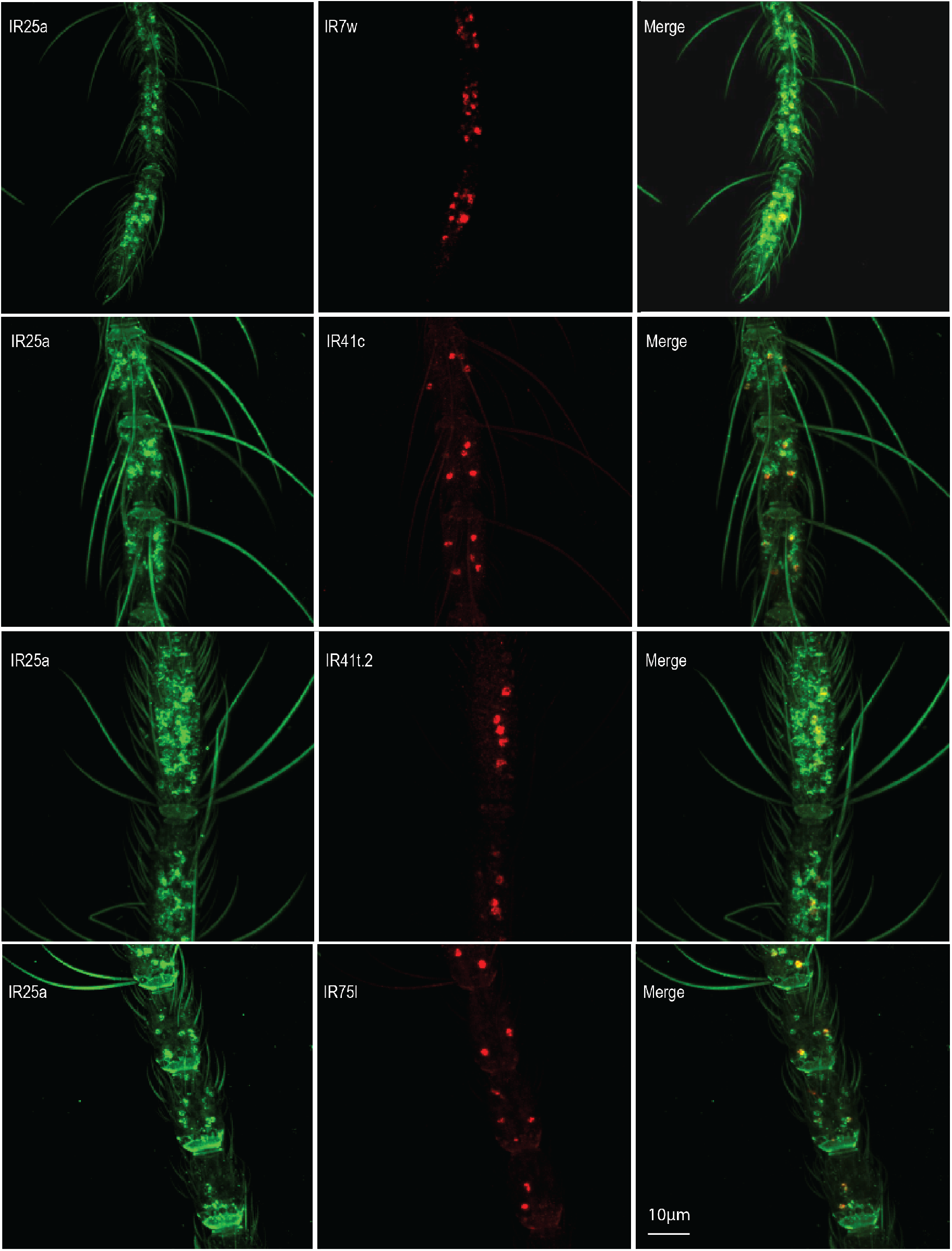
Co-expression of highly expressed tuning IRs with IR25a. In situ images showing co-expression of highly expressed tuning IRs with IR25a across the flagellomeres. Tuning IR probes were conjugated to the Alexa 647 fluorophore (red) while the IR25a co-receptor probe was linked to the Alexa 488 fluorophore (green).

**Fig 5 suppl 3:**
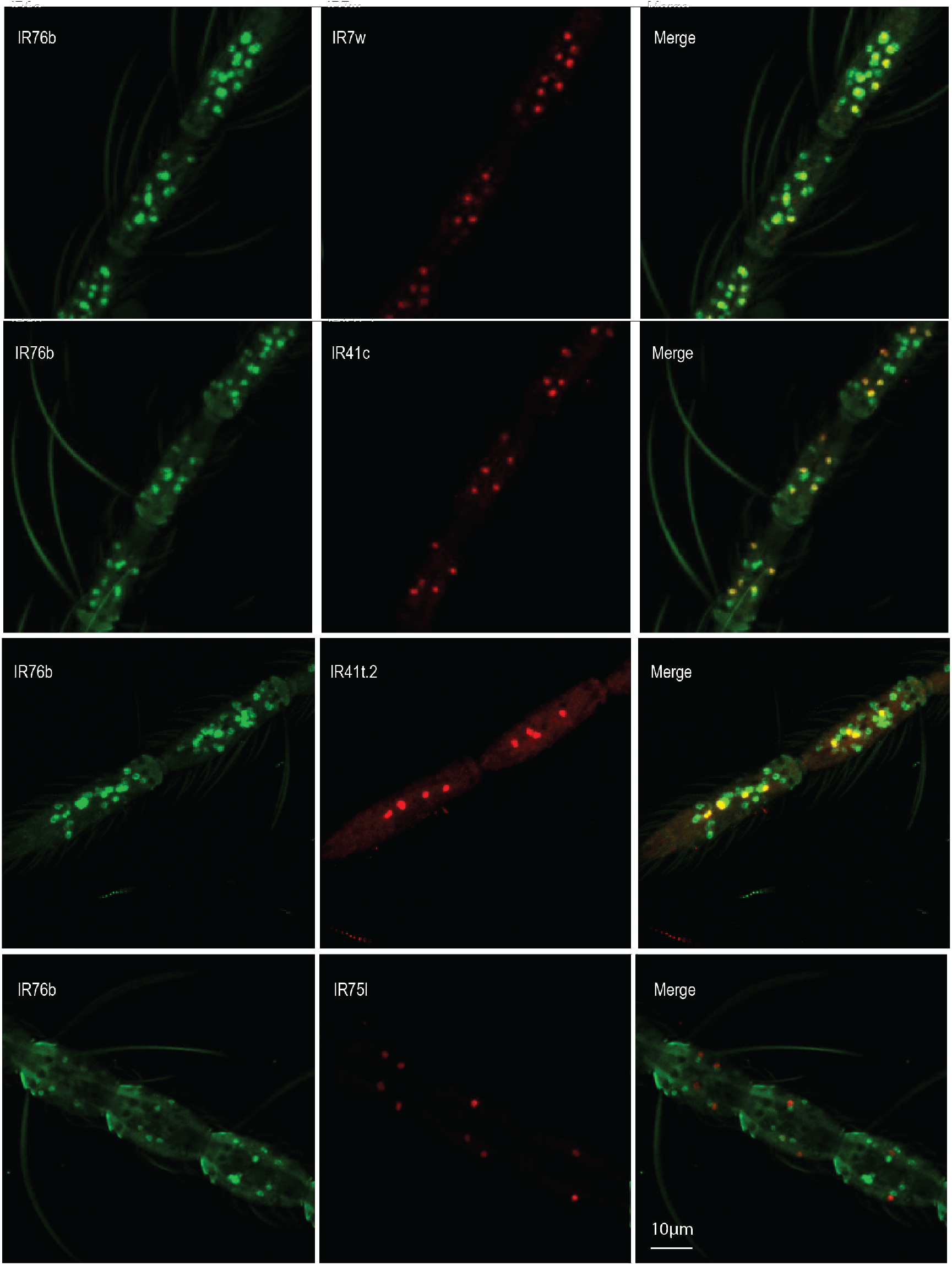
Co-expression of highly expressed tuning IRs with IR76b. In situ images showing co-expression of highly expressed tuning IRs with IR76b across the flagellomeres. Tuning IR probes were conjugated to the Alexa 647 fluorophore (red) while the IR76b co-receptor probe was linked to the Alexa 488 fluorophore (green).

